# Predicting MmpR-based bedaquiline resistance using sequence- and structure-based features

**DOI:** 10.1101/2022.12.19.521152

**Authors:** Emmanuel Rivière, Mahdi Safarpour, Pieter Meysman, Bart Cuypers, Conor Meehan, Kris Laukens, Annelies van Rie

## Abstract

Accurate molecular detection of resistance to bedaquiline, a core drug for the treatment of drug resistant tuberculosis, remains challenging. In this study, we investigated whether three sequence- and nine structure-based features describing the impact of *Rv0678* variants on the MmpR transcriptional repressor could predict the bedaquiline phenotype of isolates containing *Rv0678* variants. Paired genotypic and phenotypic data was used to train a binary random forest classifier. The mean value of the individual features was similar for resistant and susceptible variants (p≥0.05). Leave one out cross validation showed predictability of bedaquiline resistance from *Rv0678* variants when using a binary classifier with a combination of sequence and structural features (ROC AUC = 0.766; f1 score = 0.746), but the performance was too low for clinically use. Evolutionary conservation of the affected residue was the most important individual feature (mean decrease in impurity = 0.226 ± 0.003) to discriminate bedaquiline resistance from susceptibility. Prediction of bedaquiline resistance was only possible when restricting the data to missense variants, as the selected features could not be applied to *Rv0678* insertions and deletions. Additionally, prediction of bedaquiline resistance was only possible when restricting the data to isolates for which the phenotype was determined using Mycobacterial Growth Indicator Tubes, suggesting possible misclassification of the bedaquiline phenotype by other methods. The results of this study suggest that structural features describing *Rv0678* missense variants could be used to predict bedaquiline resistance, albeit not yet at the performance level required for clinical practice.

**Author summary:** Guidelines by the World Health Organization indicate that antibiotic resistant tuberculosis should be treated with bedaquiline, a newly discovered antibiotic. However, resistance to bedaquiline has already emerged and is spreading rapidly. More than 500 different variants have been discovered in the bedaquiline resistance genes in Mycobacterium tuberculosis. It is not known which DNA variants specifically drive bedaquiline resistance due to insufficiency of data. Here, we used a machine learning technique to predict whether a variant in the *Rv0678* gene confers bedaquiline resistance. To this end, for each variant we defined features that describe the impact of the variant on the MmpR (encoded by the *Rv0678* gene) protein structure. These features were then fed into a classification algorithm to predict bedaquiline resistance. We successfully developed a bedaquiline resistance prediction model, although performance was limited. We also found that accurate predictions were only possible when restricting the data to samples that were tested on the MGIT platform, a World Health Organization endorsed method to test for bedaquiline resistance. Our study shows that predicting bedaquiline resistance from protein structural features is feasible. Furthermore, our study provides new insights into phenotypic testing platforms to assess bedaquiline resistance.

## Introduction

Tuberculosis (TB), caused by members of the *Mycobacterium tuberculosis* complex (MTBC), remains one of the most challenging and deadly diseases worldwide. In 2020, 5.8 million cases of active TB and 1.5 million TB deaths were recorded[1]. While impressive progress was made during the past decades, the emergence and spread of drug resistant TB (DR-TB) threatens the goals of the End TB Strategy[2]. In 2012, bedaquiline (BDQ) became the first new antibiotic approved for treatment of DR-TB in 40 years[3]. Unfortunately, cases of BDQ treatment failure and resistance were already reported soon after its introduction[4–7].

Six genes have been listed as candidate BDQ resistance genes: *atpE, Rv0678, Rv1979c, pepQ, mmpL5*, and *mmp*S5[8]. While variants in the *atpE* gene, encoding the BDQ drug target ATP Synthase, can greatly impact the BDQ phenotype *in vitro*, these variants are rarely observed in clinical isolates[7, 9]. Clinical BDQ resistance is mostly attributed to variants in the *Rv0678* gene, which encodes a transcriptional repressor MmpR of the *mmpS5/mmpL5* efflux pump genes[10]. Loss of function variants in the *Rv0678* gene can result in upregulation of the MmpS5/MmpL5 efflux pump, of which BDQ is a substrate, with varying phenotypic effect depending on the variant[4]. Despite an understanding of the molecular mechanisms that underlie BDQ resistance, not much is known regarding specific genomic markers of BDQ resistance or susceptibility. In the 2021 WHO mutation catalogue of MTBC, only four of the 537 variants observed in the BDQ candidate resistance genes could be statistically associated with BDQ susceptibility and none with resistance[8]. A similar observation was made in a systematic review on the association of variants with phenotypic BDQ resistance[9].

This gap in knowledge regarding BDQ resistance markers limits our ability to timely diagnose BDQ resistance. Alternative methods to the standardized statistical approach to identify BDQ resistance markers should thus be explored. For pyrazinamide (PZA), another anti-tuberculosis drug, it was shown that machine learning models can accurately predict clinical PZA resistance based on sequence and structural features describing missense variants in the *pncA* gene[11]. For BDQ, a study recently elucidated the structural implications of variants in the *Rv0678* gene that drive BDQ resistance suggesting that *Rv0678* variants may cause a loss of function of the MmpR protein by disrupting protein stability, dimer interaction[12]. One study showed that a multilayer perceptron neural network could be accurately trained to predict BDQ resistance based on structural features describing variants in the *atpE* gene[13], but such analysis has not yet been performed for *Rv0678* variants. Building on these recent studies, we investigated whether a supervised machine learning approach of structural features could be used to predict clinical BDQ resistance from missense variants in the *Rv0678* gene.

## Results

### Genotypic and phenotypic data

To build a classification model which predicts BDQ resistance based on a set of three sequence- and nine structure-based features describing *Rv0678* variants, publicly available data of an individual isolate systematic review was used[9]. Of the 740 isolates in the individual isolate systematic review, 461 isolates were considered eligible for inclusion (Fig. 1). Of the 461 isolates, 420 (91.1%; 420/461) were clinical and 41 (8.9%; 41/461) were non-clinical isolates. In these 461 isolates, 197 unique genomic variants in the *Rv0678* were reported, corresponding to 194 different variants on protein-level. Of the 197 unique genomic variants, 131 (66.5%; 131/197) were missense variants, 58 (29.4%; 58/197) were indels, and eight (4.1%; 8/197) were nonsense variants. For these 461 isolates, the minimal inhibitory concentration (MIC) was determined on the Mycobacterial Growth Indicator Tube (MGIT) platform for 173 (37.5%; 173/461) isolates, on the Thermo Fisher plate platform for 178 (38.6%; 178/461) isolates, on the 7H11 platform for 172 (37.3%; 172/461) isolates, on the microplate alamar blue assay (MABA) platform for 29 (6.3%; 29/461) isolates, and on the resazurin microtiter plate assay (REMA) platform for 44 (9.5%; 44/461) isolates. For the 197 *Rv0678* variants present in the 461 isolates, 103 variants (52.3%; 103/197) were classified as ‘BDQ-resistant’ as they occurred solely or predominantly in phenotypically BDQ resistant isolates and 79 (40.1%; 79/197) were classified as ‘BDQ-susceptible’ as they occurred solely or predominantly in phenotypically BDQ susceptible isolates. The remaining 15 variants (7.6%; 15/197) could not be classified as they were observed an equal number of times in resistant and susceptible isolates.

**Fig. 1.**
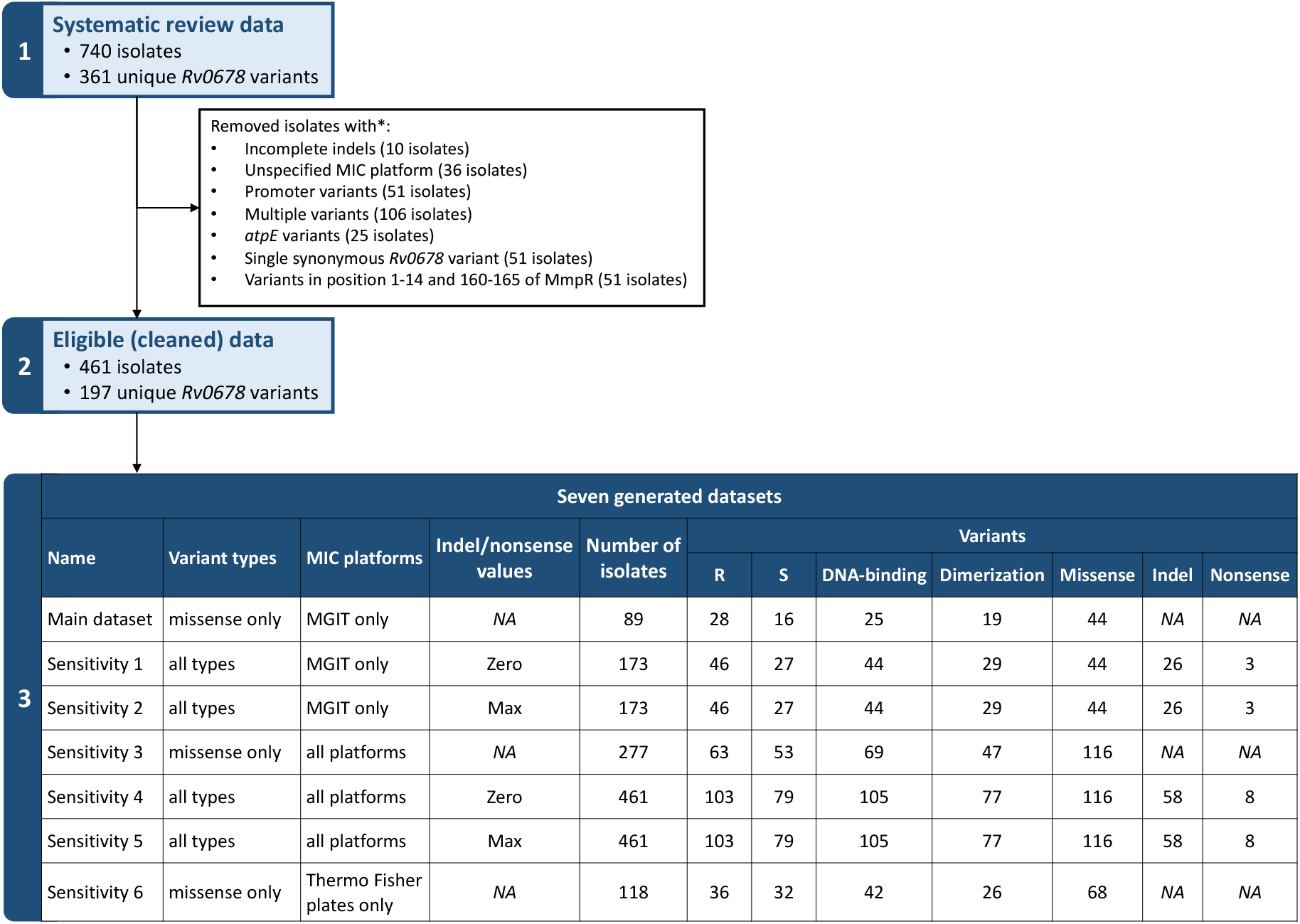
Criteria and characteristics of the different datasets. All variant characteristics were calculated after removing variants occurring an equal number of times in susceptible and resistant isolates. * These criteria are not mutually exclusive R: Resistant; S: Susceptible; MIC: Minimal Inhibitory Concentration

The main dataset of 89 isolates contained 44 unique genomic variants in the *Rv0678* gene. Of the 44 variants included in the main dataset, 16 (36.4%; 16/44) were classified as ‘BDQ-susceptible’ as they occurred solely or predominantly phenotypically BDQ susceptible isolates and 28 (63.6%; 28/44) as ‘BDQ-resistant’ as they occurred solely or predominantly phenotypically BDQ resistant isolates. Twenty-five variants (56.8%; 25/44) were located in the DNA-binding region of the MmpR transcriptional regulator and 19 variants (43.2%; 19/44) were located in the dimerization interface of the dimer (Fig. 2). When mapping the variants to both MmpR structural conformations, no obvious clustering could be observed by visual inspection.

**Fig. 2.**
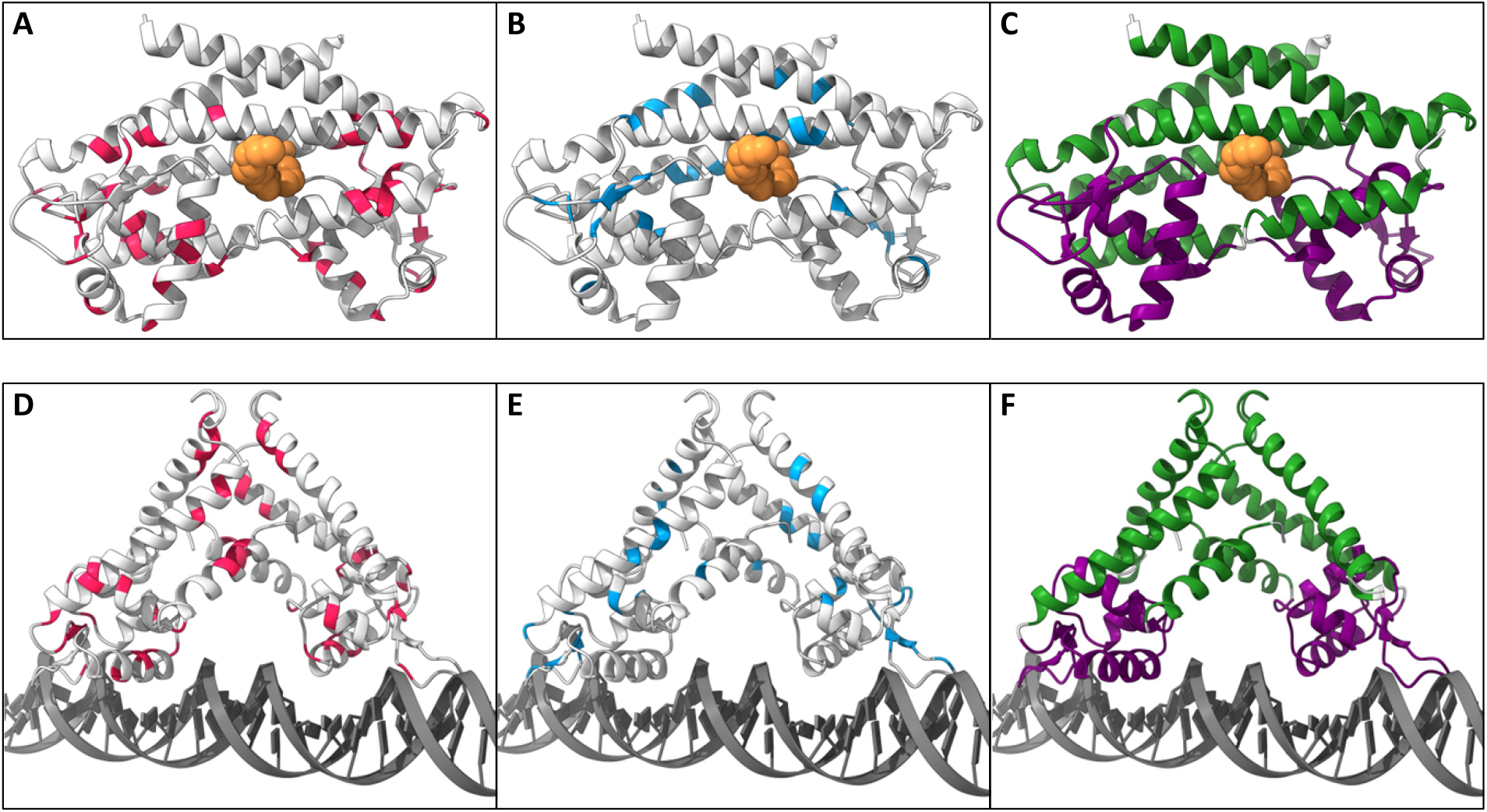
Missense variants from isolates with MGIT MIC mapped on the MmpR structure. (A) BDQ resistant (red) variants mapped on the MmpR dimer in its ligand-bound conformation. The orange molecule represents the stabilizing fatty acid ligand. (B) BDQ susceptible (blue) variants mapped on the MmpR dimer in its ligand-bound conformation. The orange molecule represents the stabilizing fatty acid ligand. (C) MmpR dimer in its ligand-bound conformation with highlighted DNA-binding domain (purple) and dimerization domain (green). The orange molecule represents the stabilizing fatty acid ligand. (D) BDQ resistant (red) variants mapped on the MmpR dimer in its DNA-bound conformation. (E) BDQ susceptible (blue) variants mapped on the MmpR dimer in its DNA-bound conformation. (F) MmpR dimer in DNA-bound conformation with highlighted DNA-binding domain (purple) and dimerization domain (green).

### Structural feature comparison

We first investigated whether any of the 12 individual features differed between *Rv0678* variants classified as ‘BDQ resistant’ and those classified as ‘BDQ susceptible’ In the main analysis (using the main dataset of 44 unique variants), the sequence-based features for *Rv0678* variants that were classified as ‘BDQ resistant’ were not statistically different from those of ‘BDQ susceptible’ variants (Mann-Whitney *U:* ConSurf p=0.096, Grantham score p=0.474, and BLOSUM62 p=0.446). Similarly, there were no differences in structure-based features between ‘BDQ resistant’ and ‘BDQ susceptible’ *Rv0678* variants [p ≥ 0.6 for mCSM-Stability (DNA and ligand), Dynamut (DNA and ligand), mCSM protein-protein interaction (DNA and ligand), mCSM-PPI RSA (DNA and ligand), and mCSM-DNA (DNA only]) (Fig. 3). Furthermore, no statistically significant differences in features were observed between variants located in the DNA-binding domain and variants located in the dimerization domain (Supplementary figures S1, S2, and S3). In a sensitivity analysis using the full dataset of 197 variants, there were also no significant differences between ‘BDQ resistant’ and ‘BDQ susceptible’ *Rv0678* missense variants (supplementary figure S4).

**Fig. 3.**
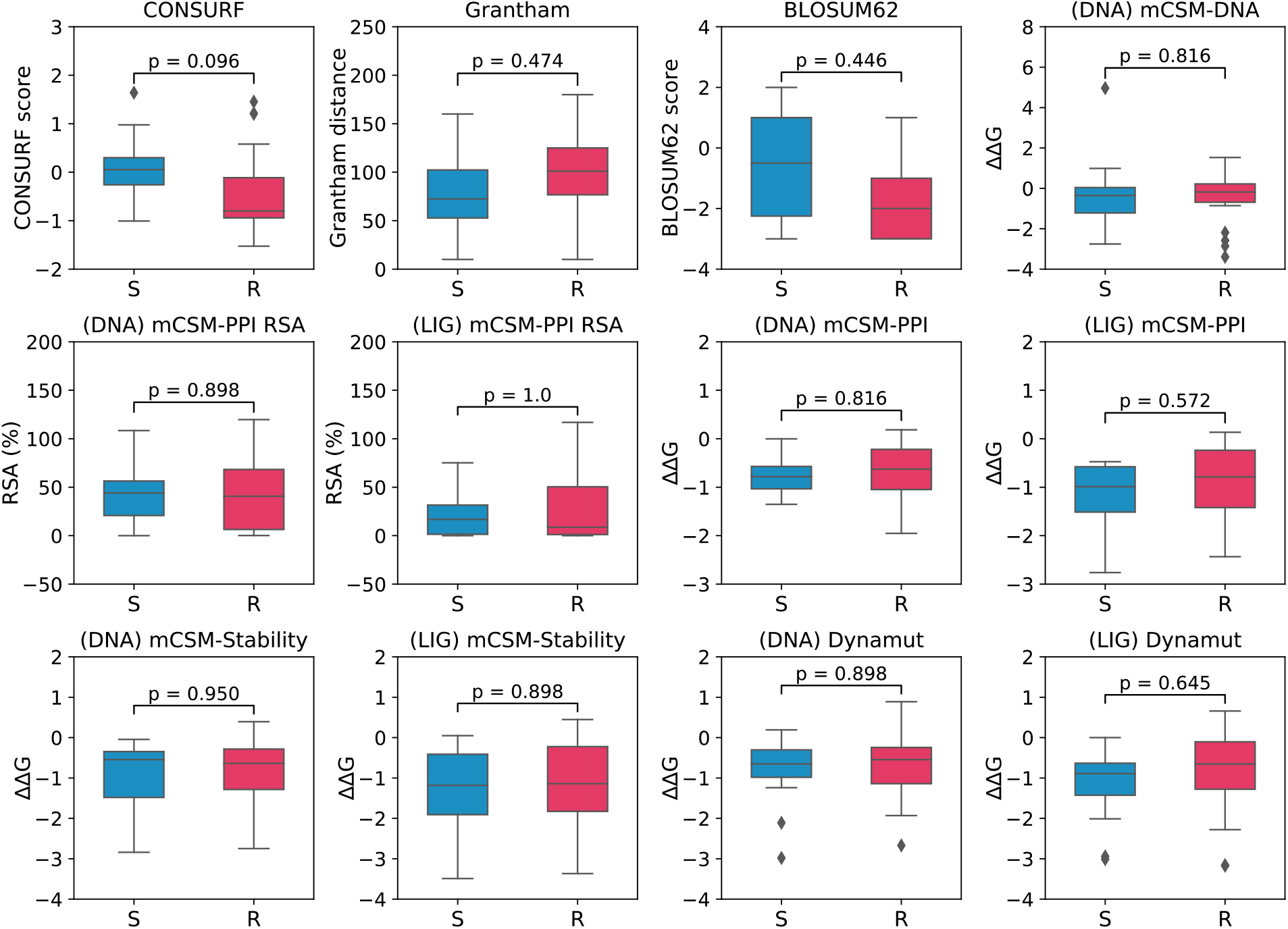
Boxplots of all calculated features for susceptible (blue) and resistant (red) missense variants in the *Rv0678* gene. The brackets indicate which conformation of the MmpR protein was used to calculate the feature (LIG = ligand-bound; DNA = DNA-bound).

### Random forest classification to predict bedaquiline resistance

To investigate whether the combined set of three sequence-based and nine structural-based features describing the functional impact of *Rv0678* variants can be used to predict phenotypic BDQ resistance, a binary random forest classification approach was used. Since the main dataset was small (n=44 observations), we opted for a leave-one-out cross-validation (LOOCV) approach to test and validate our model. Aggregating the performance over the different cross validation folds, showed that our random forest approach could predict BDQ resistance with a receiver-operator characteristic (ROC) curve area under the curve (AUC) of 0.766, precision-recall (PR) curve AUC of 0.878, and f1 score of 0.746 (Fig. 4). The feature importance analysis showed that the evolutionary conservation feature (CONSURF) is the most discriminatory feature (average importance of 0.226 ± 0.003), followed by the predicted impact on multimer formation using the ligand-bound conformation of MmpR ([LIG] mCSM-PPI, average importance of 0.104 ± 0.002). All other sequence- and structure-based features had a low (<0.1) importance in discriminating resistance from susceptible variants.

**Fig. 4.**
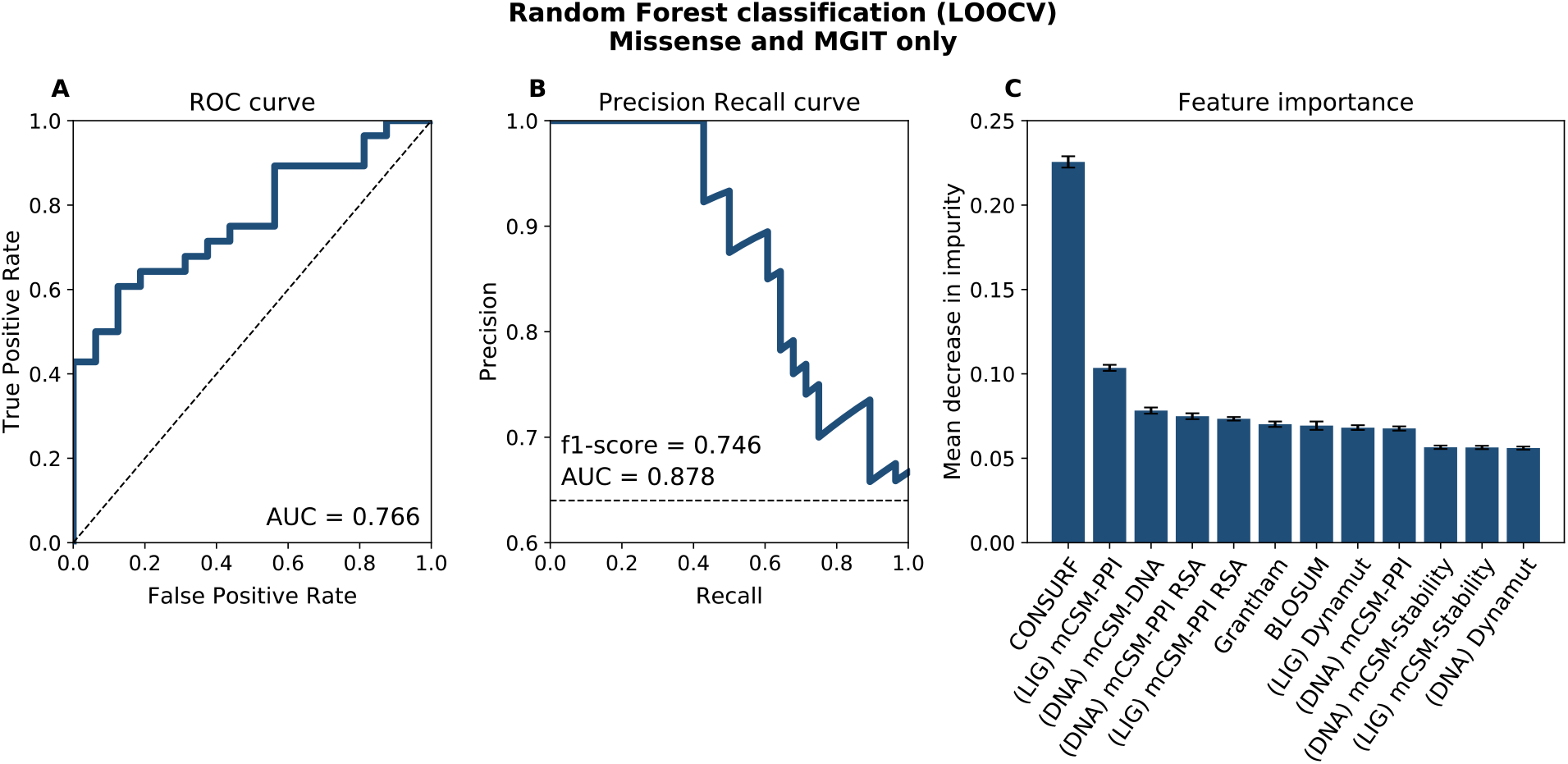
Binary classification performance using leave one out cross validation. Only missense variants and data from isolates for which a MGIT MIC was available were included. (A) Receiver Operator Characteristic curve after aggregating performance over all folds. The diagonal dotted line represents a random result. (B) Precision Recall curve after aggregating performance over all folds. The horizontal dotted line represents the ratio of positive (resistant) variants in the data set (63.6%). (C) Sorted mean decrease in impurity of all features included to build the model. The brackets indicate which conformation of the MmpR protein was used to calculate the feature (LIG = ligand-bound; DNA = DNA-bound).

In a sensitivity analysis we first used sensitivity datasets 1 and 2 (Fig. 1) to investigate whether BDQ resistance could be predicted from any type of *Rv0678* variant other than a synonymous variant, i.e. missense, indel and nonsense variants. Two analyses were performed, one where the feature values for indels were set to zero (Sensitivity dataset 1, Supplementary figure S5) and one where the feature values for indel were set to the maximum value observed in missense variants (Sensitivity dataset 2, Supplementary figure S6). These sensitivity analyses were performed with ten-fold cross validation. Both analyses resulted in low ROC AUC (0.48 and 0.53, respectively). Additional sensitivity analyses were performed using sensitivity datasets 3,4, and 5 (Fig. 1) where all phenotypic data (MGIT, Thermo Fisher plates, 7H11, MABA, and REMA) were included with and without *Rv0678* indel and nonsense variants, to investigate whether these additional data would improve BDQ resistance predictions (Supplementary figures S7, S8, S9). All these sensitivity analyses showed a close to random performance with ROC AUC ranging between 0.49 and 0.59). Finally, we performed a sensitivity analysis using dataset 6 (Fig. 1) to investigate whether BDQ resistance could be predicted from *Rv0678* missense variants reported in isolates for which the BDQ MIC was determined using the Thermo Fisher microplate assay. In contrast to the main analysis, where the BDQ phenotype was determined by MGIT, structural features could not predict the BDQ phenotype as determined by the Thermo Fisher microtiter plate DST (ROC AUC = 0.55; Supplementary figure S10).

## Discussion

The WHO recommends inclusion of BDQ in any all-oral regimen for treatment of RR-TB. Because recent studies suggest that the majority of BDQ treatment failure can be attributed to the emergence of variants in the *Rv0678* gene[4], there is an interest in the development of genomic drug susceptibility tests. Unfortunately, *Rv0678* variants occur scattered across the entire gene, without any specific hotspots of resistant variants[9, 14], are reported in both BDQ resistant and susceptible isolates[9, 15], and occur even in patients who never received BDQ treatment[16–18]. Next generation sequencing approaches to detect these resistance conferring variants thus remain constrained by our limited knowledge regarding the variants that drive BDQ resistance[19, 20]. Alternative methods are needed to discriminate BDQ resistance mediating from ‘benign’ variants.

Protein structure modelling coupled with machine learning has been applied successfully to predict resistance to TB drugs, including resistance to rifampicin by *rpoB* variants[21, 22], and isoniazid by *katG* variants[23]. Recently, a multi-layer perceptron neural network model was designed to predict pyrazinamide resistance from variants in the *pncA* gene, based on a suite of sequence- and structure-based features[11]. Interestingly, the combination of available genotypic data on PZA resistance and features describing the structural impact of variants in the *pncA* gene increased the proportion of strains for which the PZA phenotype could be correctly classified from 75% to 90%[11]. To date, only one study[13] has used structural input features to predict BDQ resistance but the analysis was restricted to *atpE* variants which are known to be of very limited clinical relevance.

We used the largest publicly available dataset of variants in the *Rv0678* gene to investigate whether three sequence-based and nine structure-based features describing the impact of these variants could be used to predict BDQ resistance. We found no statistically significant differences for any one of the 12 individual features between *Rv0678* variants occurring solely or predominantly in phenotypically resistant versus susceptible isolates, suggesting that a single feature is unlikely to be sufficient to discriminate a BDQ resistant from susceptible phenotype. This is in contrast to the findings of an *in vitro* evolution study which observed a significant difference in MmpR protein destabilization score ([*DNA*] *mCSM-Stability* and [*LIG*] *mCSM-Stability*) between missense *Rv0678* variants in BDQ resistant and susceptible isolates[12]. We could not confirm this finding as the mean scores for (*DNA*) *mCSM-Stability* and (*LIG*) *mCSM-Stability* were similar between the BDQ resistant and susceptible isolates included in our analyses. Only when combining all 12 features could a random forest classification approach predict BDQ resistance of missense variants in the *Rv0678* gene with an ROC AUC of 0.766. These results suggest that a combination of structural features could be useful to identify BDQ resistance mediating *Rv0678* variants even though the performance is currently too low for clinically use. To improve the performance of feature-based predictions of BDQ resistance, analyses on larger standardized genotypic and phenotypic datasets, inclusion of more features, and a more comprehensive approach expanding the analysis to all types of variants will be needed.

Notably, evolutionary conservation calculated using ConSurf was the best individual feature to discriminate BDQ resistance from susceptibility. This is not surprising, as conserved residues tend to be located in protein core regions and are, in general, more important to protein stability and function compared to less conserved residues[24–26]. For example, it has been previously reported that isoniazid resistance conferring variants tend to affect highly conserved residues in the *katG* gene, while variants affecting less conserved residues are associated with a lower level of isoniazid resistance[27].

Surprisingly, the BDQ resistance predictability observed for classifying missense variants was lost when including indel and nonsense variants. Due to their highly disruptive nature, indel and nonsense variants are associated with a higher fitness cost relative to missense variants[28]. For example, indels and nonsense variants have been described to confer high-level isoniazid and PZA resistance[29, 30]. In contrast, *Rv0678* indel and nonsense variants are frequently reported in clinical isolates[31, 32], even in BDQ susceptible clinical isolates[9]. The poor predictability was likely due to the difficulty in describing the impact of indel and nonsense variants with features that accurately capture the information loss on protein level[33].

Another surprising observation was that the predictability was lost when phenotypic data obtained by platforms other than the MGIT platform were included in the analysis. Although many different phenotypic platforms are used to determine BDQ resistance, a recent study showed that the MGIT platform is the most reproducible method to determine BDQ resistance, followed by broth microdilution using dry plates, such as the Thermo Fisher plates[34]. Currently, only MGIT and 7H11 are endorsed by the WHO with interim critical concentrations of 1μg/ml and 0.25μg/ml, respectively[35]. These interim critical concentrations are often based on a consensus in the absence of solid scientific data and could thus result in misclassification of resistance[36]. The loss of predictability when including MIC data obtained by any DST platform or when limiting the data to the Thermo Fisher microtiter plate assay was likely due to the misclassification of isolates as susceptible or resistant by the phenotypic methods other than the gold standard MIGT method.

Several limitations of this study should be considered. First, the data used in the analysis was derived from multiple studies with different study designs and different methodologies for phenotypic assessment which could have resulted in misclassifications. We tried to overcome this by limiting the main analysis to studies that used the gold standard MGIT platform. Second, while we excluded all isolates containing a variant in the *atpE* gene, the other candidate BDQ resistance genes (*pepQ, mmpL5, mmpS5* and *Rv1979c*) were not considered in the analysis. Although rare, variants in the *mmpL5* gene have been shown to override *Rv0678*-mediated BDQ resistance, resulting in BDQ hypersusceptibility[37]. Similarly, variants in *pepQ* and *Rv1979c* have been observed in BDQ resistant isolates, although evidence for their role in clinical BDQ resistance is very scarce[38, 39]. Third, many isolates with variants had to be excluded. We tried to overcome this by performing sensitivity analyses that included indels and nonsense variants, but isolates containing multiple variants and variants in the promotor region of *Rv0678* had to be excluded from all analyses. Finally, we could only identify 12 relevant sequence- and structure-based features. The publication of the deep learning based AlphaFold 2 has revolutionized protein structure prediction[40]. Although not yet recommended to predict the impact of variants on protein stability and function, a recent study has hinted that AlphaFold 2 could in future be used to study the effect of missense variants on protein structures[41, 42]. It will be interesting to see if improvements of the AlphaFold 2 model will enable to study of the effect of multiple variants (missense, nonsense, or indel) from *de novo* predicted protein structures compared to the wild type structures. Lastly, the clinical performance of this computational prediction model would ideally be validated using an independent dataset of clinical isolates with known BDQ phenotype and *Rv0678* variants. Since BDQ is still a relatively novel drug, such datasets were not available at the time this study was conducted.

Many studies have shown that PZA resistance-conferring and benign variants are frequently observed across the entire length of the *pncA* gene without any apparent resistance hotspots, resulting in a markedly low sensitivity to identify PZA resistance when only considering the genotypic information[43, 44]. Similarly, *Rv0678* variants occur across the entire gene and the standard statistical approach is inadequate to accurately diagnose BDQ resistance from genotypic data[9, 44]. Our study suggest that a combination of sequence- and structure-based features may, in future, be able to predict BDQ resistance from *Rv0678* missense variants. This could facilitate the development of sequencing-based strategies as diagnostics to detect BDQ resistance and prevent emergence and spread of BDQ-resistant TB.

## Material and Methods

### Data collection and preparation

Publicly available data from an individual isolate systematic review and meta-analysis were used for model development[9]. The systematic review data contains all studies that reported on genotypic and phenotypic BDQ resistance for both clinical and non-clinical *Mycobacterium tuberculosis* (*Mtb*) isolates published between January 2008 and December 2021. For this analysis, isolates were classified as resistant or susceptible using the drug susceptibility testing (DST) platform-specific minimal inhibitory concentration (MIC) cut-off values of 1 μg/ml for Mycobacterial Growth Indicator Tubes (MGIT), 0.25 μg/ml for 7H11, and 0.125 μg/ml for the microplate alamar blue assay (MABA), resazurin microtiter plate assay (REMA), and the Thermo Fisher microplate assay. When MIC values were reported for the same isolate on different platforms, then the MIC result obtained by MGIT was prioritised and selected, followed by the Thermo Fisher plates, and then 7H11[34]. *Rv0678* variants that were reported in more than one isolate were classified as either resistant or susceptible based on whether the majority of isolates containing the variant were phenotypically BDQ resistant or susceptible.

Several isolates were excluded from the analysis (Fig. 1). First, isolates which contained a variant in the *Rv0678* promoter region, or more than one missense variant in the *Rv0678* coding region were excluded because features could not be calculated for these isolates. Second, isolates with a variant in both the *atpE* and *Rv0678* gene were also excluded because the presence of these *atpE* variants could confound the genetic causality of BDQ resistance by the *Rv0678* variant. When information on the *atpE* gene was missing, the isolate was considered *atpE wild type* and included in the analysis. Third, isolates for which the MIC platform used was not specified were excluded because this information is needed to classify the isolate as phenotypically resistant or susceptible. Fourth, variants that occurred an equal number of times in resistant and susceptible isolates could not be classified as resistant or susceptible. Isolates containing these variants were excluded. Fifth, all isolates with a variant affecting amino acid positions 1-14 and 160-165 of the MmpR protein were excluded because these regions are not included in the MmpR crystal structure, thus precluding the assessment of the impact of the *Rv0678* variant on the protein. Finally, all synonymous variants in the *Rv0678* gene were ignored for the analysis.

For the main analysis, all isolates containing an indel variant or missense variant resulting in a premature stop-codon (nonsense) were excluded, and only isolates for which the MIC was determined on the MGIT platform, the current gold standard, were included in the analysis. Five sensitivity analysis datasets were constructed to investigate the impact of inclusion of non-missense variants and isolates for which the MIC was determined using non-MGIT platforms (Fig. 1).

### Feature extraction

For each missense variant, the evolutionary conservation score of the affected MmpR residue was calculated with ConSurf (version 2016) using the default parameters with the *Mtb* MmpR crystal structure (PDB ID: 4NB5) as a guide structure[45, 46]. The Grantham score was calculated to estimate the distance between each replacing amino acid pair[47]. A BLOSUM62 substitution matrix was used to estimate the level of conservation of each MmpR amino acid replacement[48].

To calculate the structural features of each missense variants, two MmpR structural conformations were used to represent both the ligand-bound state and the DNA-bound state. For the ligand-bound state, the experimentally determined crystal structure of *Mtb* MmpR was used (PDB ID: 4NB5)[46]. For the DNA-bound state, a published structure, determined through homology modelling of the DNA-binding domain of MmpR based on other *Mtb* MarR-like transcriptional regulators, was used[12]. The effect of the missense variants on both states of the MmpR protein was assessed with a suite of established and publicly available mCSM tools (http://biosig.unimelb.edu.au/biosig/tools). The effect on protein structure was assessed by mCSM-Stability[49] and Dynamut2[50]. Changes in protein-protein affinity and the relative solvent accessibility (RSA) of the MmpR dimer in both structural conformations was predicted with mCSM-PPI2[51]. The impact on the DNA binding affinity of the MmpR transcriptional regulator was predicted with mCSM-DNA for MmpR in its DNA-bound state only[52].

For *Rv0678* indels and nonsense variants, ConSurf scores were calculated by averaging the evolutionary scores of the residues in the affected regions. Because the other features could not be calculated for indel and nonsense variants, binary features indicating the variant type and whether the variant causes a disruption of the DNA reading frame were created for all variants. For each indel, two sets of feature scores were created. In one set the feature scores for all indel and nonsense variants were set to zero to investigate whether the binary features describing the variant type are sufficient to predict BDQ resistance. In the other set, the feature scores for all indel and nonsense variants were set to the maximum score of the missense variants, to accurately represent the highly disruptive nature of such variants.

The MmpR structures in ligand-bound and DNA-bound conformations were visualised in ChimeraX and the dimerization and DNA-binding domains were defined as described by *Radhakrishnan et al.[46, 53]* Feature distributions were visualized using the Matplotlib library (version 3.1.2) in Python[54] and were compared between resistance and susceptibility variants using the Mann-Whitney *U* test. P-values were adjusted for false discovery rate using the Benjamini-Hochberg procedure and were considered significant when p-value < 0.05.

### Construction of machine learning classifier

In order to predict whether a variant would be associated with a BDQ resistant phenotype, a binary random forest classifier was trained using the Scikit-learn Python module (version 1.0.2)[55] with default parameters and the number of trees set to 1000. Classification models were trained using repeated ten-fold cross validation. Models were evaluated by calculating the receiver operator characteristic curve (ROC) with area under the curve (AUC) and precision recall curve (PR) with f1 score after aggregating performance measures over the different folds, using the Scikit-learn Python module (version 1.0.2). When the dataset was too small (less than 50 observations) for a ten-fold cross validation approach, a leave-one-out cross validation (LOOCV) approach was adopted instead. Feature importance, computed as the mean decrease of impurity within each tree, was calculated using the Scikit-learn Python module (version 1.0.2). ROC curve, PR curve and extracted feature importance were visualised using the Matplotlib package (version 3.1.2) in Python[54].

## Acknowledgements

We thank Joshua J. Carter (Stanford University and CRyPTIC consortium) for providing the DNA-bound structure of MmpR. This work was supported by Research Foundation Flanders (FWO Odysseus G0F8316N and FWO Strategic Basic Research 1S39119N).

## Supporting information captures

**S1 Figure. DNA-binding domain features.** Boxplots of all calculated features for susceptible (blue) and resistant (red) missense variants located in the *Rv0678* gene region encoding the DNA-binding domain of MmpR (DNA-binding subset of the Main dataset). The brackets indicate which conformation of the MmpR protein was used to calculate the feature (LIG = ligand-bound; DNA = DNA-bound).

**S2 Figure. Dimerization domain features**. Boxplots of all calculated features for susceptible (blue) and resistant (red) missense variants located in the *Rv0678* gene region encoding the dimerization domain of MmpR (Dimerization subset of the Main dataset). The brackets indicate which conformation of the MmpR protein was used to calculate the feature (LIG = ligand-bound; DNA = DNA-bound).

**S3 Figure. DNA-binding vs dimerization features**. Boxplots of all calculated features for missense variants located in the *Rv0678* gene region encoding the DNA-binding domain (red) and missense variants located in the *Rv0678* gene region encoding the dimerization (blue) domain of MmpR. The brackets indicate which conformation of the MmpR protein was used to calculate the feature (LIG = ligand-bound; DNA = DNA-bound).

**S4 Figure. All features on all pDST platforms**. Boxplots of all calculated features for susceptible (blue) and resistant (red) missense variants located in the *Rv0678* gene using all available data (all pDST platforms; Sensitivity dataset 3). The brackets indicate which conformation of the MmpR protein was used to calculate the feature (LIG = ligand-bound; DNA = DNA-bound).

**S5 Figure. Random forest classification using MGIT only and indel values set to zero**. Binary classification performance using 10-fold cross validation using MGIT phenotypic data only for missense and indel variants (Sensitivity dataset 1 in Figure 1). Indel feature values were set to zero. (A) Receiver Operator Characteristic curve after aggregating performance over all folds. The diagonal dotted line represents a random result. (B) Precision Recall curve after aggregating performance over all folds. (C) Sorted mean decrease in impurity of all features included to build the model. The brackets indicate which conformation of the MmpR protein was used to calculate the feature (LIG = ligand-bound; DNA = DNA-bound).

**S6 Figure. Random forest classification using MGIT only and indel values set to max.** Binary classification performance using 10-fold cross validation using MGIT phenotypic data only for missense and indel variants (Sensitivity dataset 2 in Figure 1). Indel feature values were set to the maximum value among the missense variants. (A) Receiver Operator Characteristic curve after aggregating performance over all folds. The diagonal dotted line represents a random result. (B) Precision Recall curve after aggregating performance over all folds. (C) Sorted mean decrease in impurity of all features included to build the model. The brackets indicate which conformation of the MmpR protein was used to calculate the feature (LIG = ligand-bound; DNA = DNA-bound).

**S7 Figure. Random forest classification using all data without indels**. Binary classification performance using 10-fold cross validation using all available phenotypic data for missense variants only (Sensitivity dataset 3 in Figure 1). (A) Receiver Operator Characteristic curve after aggregating performance over all folds. The diagonal dotted line represents a random result. (B) Precision Recall curve after aggregating performance over all folds. (C) Sorted mean decrease in impurity of all features included to build the model. The brackets indicate which conformation of the MmpR protein was used to calculate the feature (LIG = ligand-bound; DNA = DNA-bound).

**S8 Figure. Random forest classification using all data with indels set to zero**. Binary classification performance using 10-fold cross validation using all available phenotypic data for missense and indel variants (Sensitivity dataset 4 in Figure 1). Indel feature values were set to zero. (A) Receiver Operator Characteristic curve after aggregating performance over all folds. The diagonal dotted line represents a random result. (B) Precision Recall curve after aggregating performance over all folds. (C) Sorted mean decrease in impurity of all features included to build the model. The brackets indicate which conformation of the MmpR protein was used to calculate the feature (LIG = ligand-bound; DNA = DNA-bound).

**S9 Figure. Random forest classification using all data with indels set to max**. Binary classification performance using 10-fold cross validation using all available phenotypic data for missense and indel variants (Sensitivity dataset 5 in Figure 1). Indel feature values were set to the maximum value among the missense variants. (A) Receiver Operator Characteristic curve after aggregating performance over all folds. The diagonal dotted line represents a random result. (B) Precision Recall curve after aggregating performance over all folds. (C) Sorted mean decrease in impurity of all features included to build the model. The brackets indicate which conformation of the MmpR protein was used to calculate the feature (LIG = ligand-bound; DNA = DNA-bound).

**S10 Figure. Random forest classification using Thermo Fisher plate data only without indels**. Binary classification performance using leave one out cross validation. Only missense variants and data from isolates for which a Thermo Fisher plate MIC was available were included. (A) Receiver Operator Characteristic curve after aggregating performance over all folds. The diagonal dotted line represents a random result. (B) Precision Recall curve after aggregating performance over all folds. The horizontal dotted line represents the ratio of positive (resistant) variants in the data set (52.9%). (C) Sorted mean decrease in impurity of all features included to build the model. The brackets indicate which conformation of the MmpR protein was used to calculate the feature (LIG = ligand-bound; DNA = DNA-bound).

**S11 Appendix. Source data.**

